# Transthyretin amyloid fibrils adopt distinct folds in the brain

**DOI:** 10.64898/2026.05.12.724695

**Authors:** Shumaila Afrin, Binh An Nguyen, Parker Bassett, Maria del Carmen Fernandez-Ramirez, Rose Pedretti, Layla Villalon, Candace Kelly, Christian Lopez, Mihir Madabushi, Annie Zhou, Ines Reis, Ricardo Taipa, Bret Evers, Lorena Saelices

## Abstract

Amyloid deposition in the central nervous system is increasingly recognized in transthyretin (ATTR) amyloidosis, particularly in patients with prolonged survival following liver transplantation or disease-modifying therapies. However, the structural basis of transthyretin aggregation in the brain remains unknown. Here we determine cryo-electron microscopy (cryo-EM) structures of *ex vivo* brain-derived ATTR fibrils from patients carrying the ATTRv-V30M and ATTRv-V30G variants. Both fibrils adopt folds distinct from those previously reported in peripheral tissues and the vitreous humor. V30M fibrils exhibit a continuous ordered core spanning residues Pro11–Asn124, whereas V30G fibrils consist of a substantially reduced ordered core, revealing pronounced structural divergence even within the same tissue environment. Despite this diversity, comparative analyses identify conserved regions across ATTR fibrils, including a segment implicated in transthyretin aggregation and targeted for diagnostic and therapeutic development. These results provide direct evidence that local tissue context can shape amyloid fibril architecture in human disease.

## Introduction

ATTR amyloidosis is a systemic protein misfolding disorder driven by the aggregation of transthyretin, a protein produced predominantly by the liver but also locally within the choroid plexus and retinal epithelium^2^. More than 220 mutations have been associated with cardiomyopathy, polyneuropathy, or both^3^, highlighting the widespread and heterogeneous nature of the disease. Despite this systemic involvement, deposition within the central nervous system (CNS) has traditionally been considered rare and largely restricted to oculoleptomeningeal variants^4^. However, growing clinical and pathological evidence indicates that transthyretin also accumulates in the brain of patients carrying common mutations, including ATTRv-V30M, particularly following liver transplantation or treatment with therapies that do not target locally produced transthyretin^5,6^. In these settings, brain deposition is observed in up to 31% of neuropathic patients^7,8^, revealing a clinically significant but underrecognized site of disease. As survival improves following disease-modifying therapies, CNS amyloid deposition is emerging as an increasingly recognized manifestation of ATTR amyloidosis for which effective diagnostics and therapeutics remain lacking^9^. Despite its increasing clinical relevance, the molecular and structural basis of transthyretin aggregation in the brain remains unknown.

Recent advances in cryo-electron microscopy (cryo-EM) have enabled the determination of near-atomic resolution structures of ATTR fibrils extracted from multiple patients, tissues, and mutations, establishing a structural framework for understanding transthyretin aggregation^10-18^. Systematic structural analyses across organs and variants, including studies from our group, show that the ordered cores of fibrils from peripheral tissues and the vitreous humor of the eye are composed of N- and C-terminal fragments of similar lengths. Most of these fibrils adopt a conserved conformation that we previously defined as the closed gate fold, although local variations, particularly in the C-terminal “gate” region, have been observed, including in vitreous humor-derived fibrils^10-18^. This apparent structural conservation is maintained despite substantial clinical heterogeneity, suggesting that a common folding pathway underlies fibril formation across tissues. Whether this structural framework extends to the brain, where the local tissue context differs from both peripheral tissues and the eye, remains unknown.

To define the structural basis of ATTR amyloidosis in the brain, we determined cryo-EM structures of amyloid fibrils extracted from two patients carrying the ATTRv-V30G and ATTRv-V30M variants (hereafter referred to as V30G and V30M, respectively). We find that these fibrils adopt folds that differ from those previously reported in peripheral tissues and the vitreous humor of the eye, indicating that transthyretin aggregation is not constrained to a single conserved fold across tissues. Instead, our results reveal a previously unrecognized dimension of tissue-specific structural polymorphism in ATTR amyloidosis. These findings indicate that transthyretin fibrils formed in the brain adopt folds distinct from those previously observed in peripheral tissues and the eye, providing a framework to link fibril structure with disease phenotype. Notably, despite these differences, comparison across all known ATTR fibril structures reveals conserved regions, including a segment that we determined to drive the aggregation of transthyretin *in vitro*^19^, and has been exploited as a target for the development of diagnostic and therapeutic strategies^19-25^.

## Results

### Amyloid deposition and fibril composition in ATTR brain tissues

We analyzed amyloid deposition and fibril composition in brain tissue samples extracted from two patients with ATTR amyloidosis, one carrying the V30G variant and one carrying the V30M variant (Supplementary Table 1). The V30M carrier initially presented with polyneuropathy (34 years old) and underwent liver transplantation, later developing CNS manifestations after a prolonged disease course (29 years disease course). The V30G carrier presented with predominant CNS involvement (45 years old) without evidence of peripheral disease, with a short disease course (patient died within 3 months of diagnosis) as previously reported^26^. To confirm the presence of amyloids in these tissues, we performed histological analysis, including hematoxylin and eosin (H&E) staining to highlight tissue architecture, and Congo red staining and Thioflavin-S staining to reveal amyloid deposits (Fig. 1a). We observed cerebral amyloid angiopathy in leptomeningeal vessels, with amyloid deposits along the vessel walls, along with small amyloid deposits in the leptomeninges in close proximity to the cortical surface in V30M. In V30G, we observe a diffused deposition of amyloid in the fibrous tissue of leptomeninges, also involving the outer layers of some vessels.

**Figure 1:**
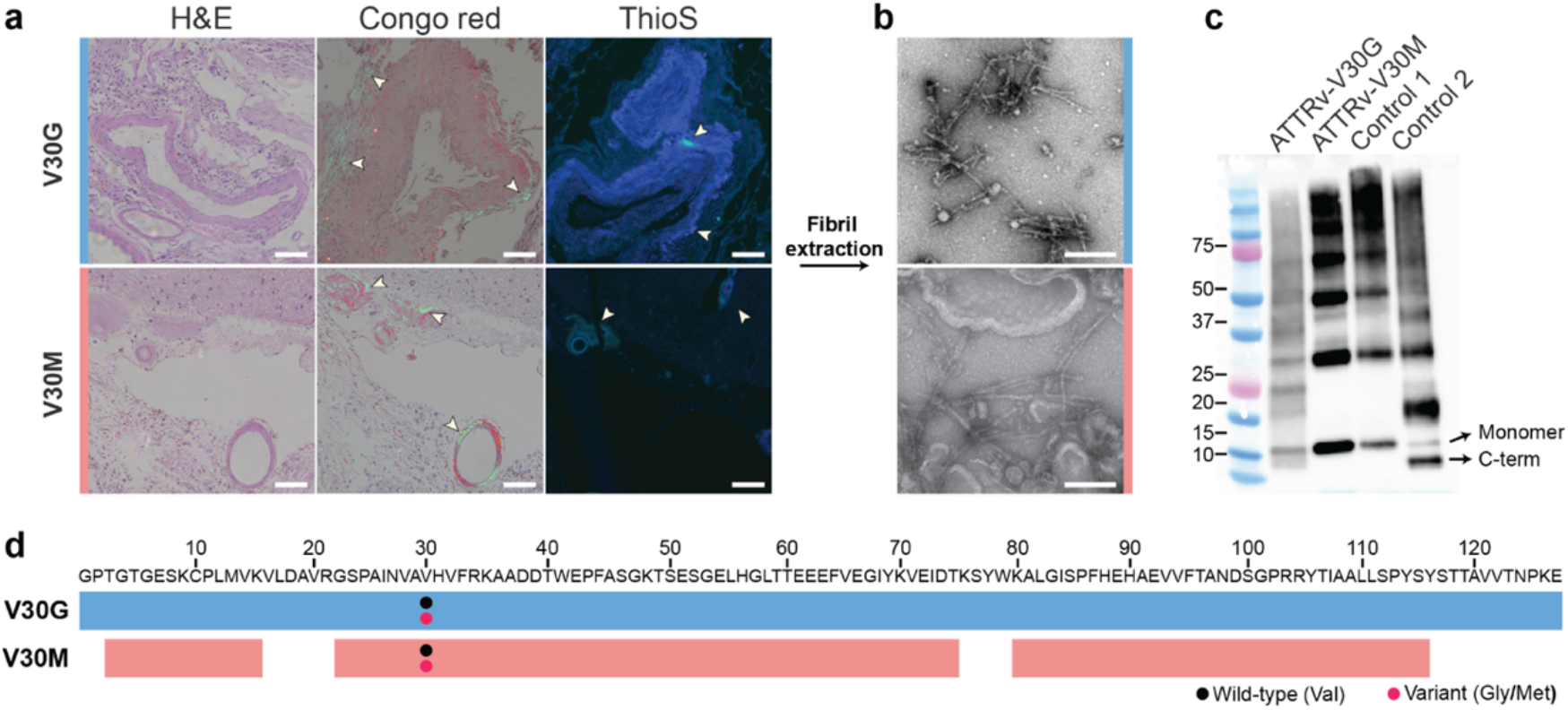
Histological and biochemical characterization of brain-derived ATTR fibrils. **a**, Histological analysis of tissue sections from the V30G (top) and V30M brain (bottom). H&E staining shows tissue architecture, Congo red staining confirms amyloid deposition, and Thioflavin S (labeled as ThioS) fluorescence highlights amyloid aggregates. Scale bars, 100 µm. **b**, Negative-stain transmission electron microscopy of fibrils following extraction from brain tissues, confirming the presence of abundant fibrillar material in both V30G (top) and V30M (bottom) samples. Scale bars, 200 nm. **c**, Western blot analysis using an antibody against a C-terminal region of transthyretin. Bands corresponding to full-length monomer and C-terminal fragments are marked with arrows. Control 1 and control 2 are cardiac V30M fibrils extracted from patients with type B and type A ATTR amyloidosis, respectively,^1^ shown for comparison. Type B and type A ATTR amyloidosis are associated with distinct fibril composition, histological distribution, and clinical manifestations^1^. **d**, Mass spectrometry sequence coverage of transthyretin in V30G and V30M fibrils. Peptide mapping shows near-complete coverage of the transthyretin sequence. Mutation sites are indicated. Peptides spanning residue 30 were detected with both wild-type valine and mutant glycine or methionine in the respective samples, confirming the presence of variant protein within the deposits and heterozygosity.

We extracted amyloid fibrils from these tissues using an optimized water-EDTA based protocol adapted from a previously described method. This gentle method solubilizes amyloid fibrils from multiple tissue types preserving their structure^10-16,27-29^. We confirmed the presence of well dispersed amyloid fibrils using transmission electron microscopy with negative staining (Fig. 1b). Immunoblotting showed the presence of transthyretin in both fibril extracts (Fig. 1c). We further validated protein identity and sequence coverage by mass spectrometry, which revealed near-complete coverage of the transthyretin sequence in fibrils from both fibril extracts (Fig. 1d). Peptides corresponding to both wild-type (Val30) and mutant (Gly30 or Met30) residues were detected, confirming the heterozygosity of these patients. We also detected signature proteins that usually co-purify with the fibrils, including serum amyloid P-component, apolipoprotein E and A-IV, heparan sulfate proteoglycans, complement components, clusterin, and vitronectin^30^.

### Cryo-EM reveals distinct structures of brain-derived ATTR fibrils

We first determined the structure of V30G fibrils extracted from brain tissues using cryo-EM imaging (Fig. 2a). Reference-free two-dimensional (2D) classification of V30G fibrils revealed a single predominant morphology consisting of one protofilament, with a helical crossover distance of 980 Å, a rise of 4.85 Å per layer, and C1 symmetry (Fig. 2b). Three-dimensional (3D) reconstruction produced a single class (Fig. 2c), and the 3.9 Å density map enabled placement of residues Ser52 to Ser100 of transthyretin into the ordered core (Fig. 2d and Supplementary Fig. 1a–b). N-terminal residues from Gly1 to Gln51 were not resolved, placing the V30G variant site outside the structured core. Although partial density was observed for the C-terminal end of the sequence beyond residue Ser100, this segment could not be reliably modeled (Fig. 2d).

**Figure 2:**
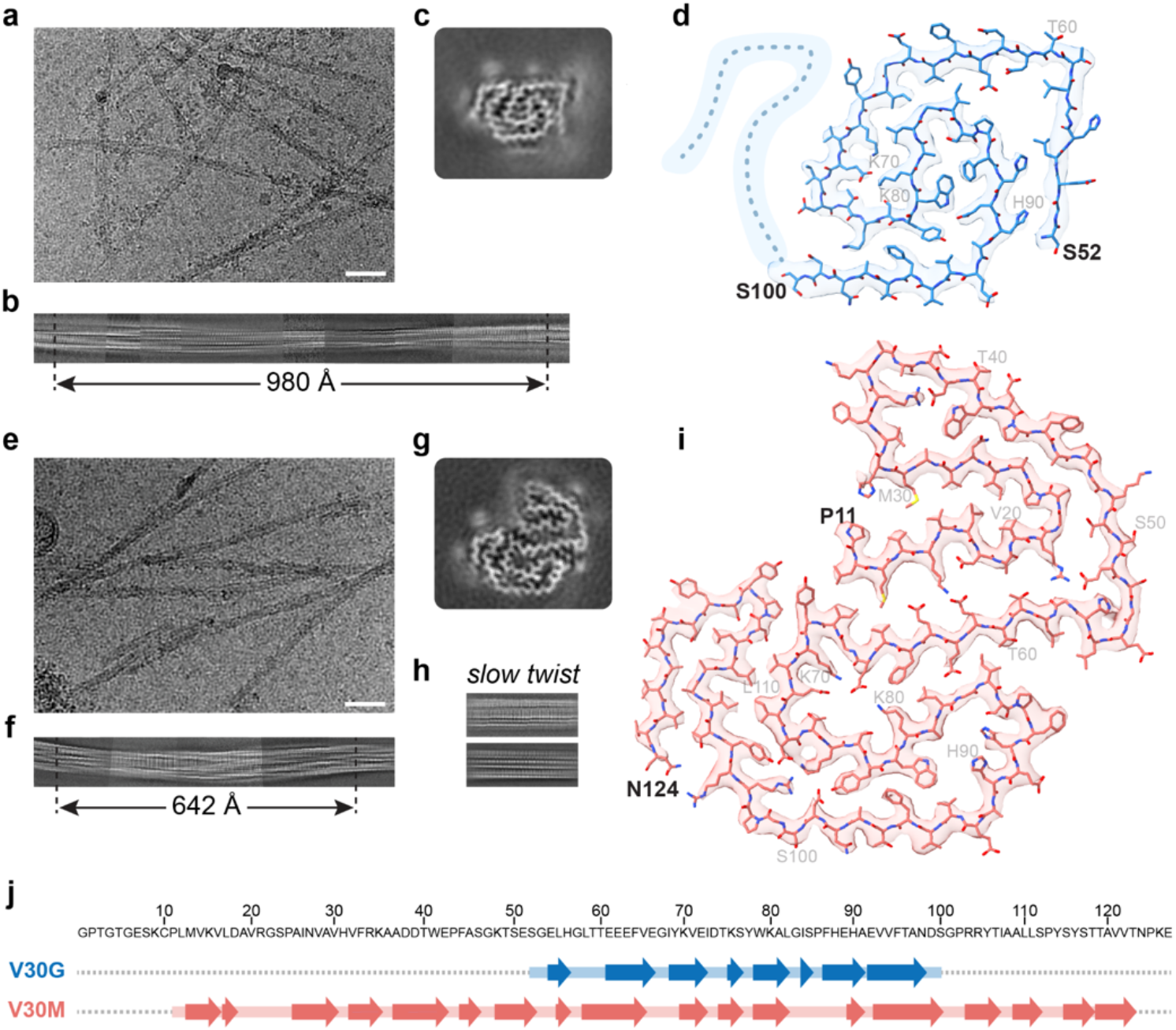
Cryo-EM structures of brain-derived V30G and V30M fibrils. **a, e**. Representative cryo-EM micrographs of V30G (a) and V30M fibrils (e). **b, f**. Reference-free 2D classes; stitched classes reconstruct one full crossover for twisted fibrils. **h**. Reference-free 2D classes for slow-twisting V30M fibrils. **c, g**. Representative 3D class averages of V30G (c) and V30M (g) fibrils. **d, i**. Top views of a single fibril layer with the atomic model fitted into the cryo-EM density for V30G (d) and V30M (i). The V30G core comprises residues Ser52– Ser100, whereas the V30M core spans residues Pro11–Asn124. Dotted lines in **d** indicate regions with visible density in the 3D class average that were not modeled due to insufficient resolution. **j**. Sequence of transthyretin with secondary structure indicated below the sequence. Corresponding β-strand assignments in V30G fibrils (blue) and V30M fibrils (salmon) are shown below, illustrating differences in secondary structure organization. Dotted lines represent regions not resolved as part of the amyloid core.

Using the same approach, we next determined the structure of V30M fibrils. 2D classification revealed a major morphology with slow or no twist and a minor twisting morphology (Fig. 1e-h). The relative abundance of these morphologies was not quantified, as particle picking was intentionally biased toward twisted fibrils. We stitched together representative 2D class averages of the twisted fibrils and observed a helical crossover distance of 642 Å (Fig. 2f). The rise was later calculated as 4.75 Å per layer with C1 symmetry. Subsequent 3D classification yielded in a single class (Fig. 2g), with a resulting density map resolved to 3.4 Å, enabling model building of residues Pro11 to Asn124 into the ordered core (Fig. 2i and Supplementary Fig. 1c-d). Although residue 30 lies within the ordered core, we could not unambiguously assign its identity as valine or methionine; for simplicity, all depictions of the V30M structure in this paper show methionine at position 30 (Fig. 2i).

### V30G and V30M fibrils exhibit distinct folds and stabilization features

The V30G fibril core comprises eight β-strands, accounting for approximately 70% of the ordered core of 48 residues (Fig. 2j). The fibril layers adopt a relatively planar arrangement, with a subunit height variation of ∼5.81 Å along the fibril axis (Supplementary Fig. 2a). Within the same layer, residues in β3, β4 and β5 further stabilize the fold through multiple potential interactions, including salt bridges between Glu72 and Lys70 and between Glu72 and Lys80, as well as hydrogen bonds between Thr75 and Glu72 and between Ser77 and Lys80 (Supplementary Fig. 2b). The fibril is further stabilized by extensive intermolecular hydrogen bonding, aromatic stacking interactions, and polar ladders (Supplementary Fig. 2c).

Additional stabilization arises from buried hydrophobic clusters. One cluster, involving Phe95, Val93, Ala91 Trp79, Phe87, Tyr78 and Ala81 residues, includes residues from the β-strand F of the native structure of transthyretin (Supplementary Fig. 3a). The second hydrophobic cluster is formed by Leu82, Ile68, Val65, Ile84, and Pro86. It is worth noting that the N-terminal segment that provides a key stabilizing dry steric zipper in all peripheral ATTR fibrils^10-16^ is missing in the V30G ordered core. Consistent with this observation, the estimated free energy of solvation for V30G fibrils is −19.8 kcal per layer (−0.40 kcal per residue), which lies on the lower end of the stability range reported for ATTR fibrils^16^, suggesting comparatively reduced packing stability (Supplementary Fig. 3b).

The V30M fibril core represents the longest ATTR fibril structure reported to date and comprises eighteen β-strands, accounting for approximately 70% of the ordered core of 113 residues (Fig. 2j). In contrast to V30G fibrils, V30M layers display a non-planar arrangement with a subunit height variation of ∼13.65 Å that enables interlayer interactions within the fibril (Supplementary Fig. 4a). The overall structure, and in particular the turns between β-strands, are extensively stabilized by a network of buried salt bridges across each layer and throughout the fibril fold (Supplementary Fig. 4b-f). Interlayer salt bridges include interactions between Arg34 of layer *n* and Asp39 of layer *n-1*, and several other interlayer electrostatic interactions (Supplementary Fig. 4b-f). Furthermore, polar ladders formed by asparagine residues extend along the fibril axis, contributing to interlayer stabilization. Within individual layers, further stabilization is provided by intralayer salt bridges (Supplementary Fig. 4b-f).

In addition to polar interactions, the structure is further stabilized by extensive hydrophobic interactions, with multiple clusters distributed throughout the fibril core. The most prominent cluster is located at the N-terminal region, where densely packed nonpolar residues form a dry steric zipper that likely represents one of the major stabilizing features of the fold (Supplementary Fig. 3c-d). This stabilizing cluster is conserved across peripheral ATTR fibril structures, but adopts a different orientation relative to the core^10-12,14-16^. Additionally, in V30M brain fibrils, the presence of a continuous transthyretin segment, rather than the two disconnected fragments observed in peripheral fibrils^10-12,14-16^, introduces additional structural elements that enable new stabilizing interactions and contribute to enhanced stability within and across fibril layers. Consistent with this, the estimated free energy of solvation is −74.7 kcal/mol per chain and −0.66 kcal per residue, placing it at the upper end of the stability range reported for ATTR fibrils^16^ (Supplementary Fig. 3d).

Unresolved densities are observed in both the V30G and V30M maps, but their identity could not be determined (Supplementary Fig. 5a-b). Although in the V30G structure there are five distinct unresolved densities, two appear to be extensions of the ordered core upstream of residue Ser52 and downstream of residue Ser100 (Supplementary Fig. 5a, i-ii). The other unresolved densities in both structures appear disconnected from the ordered core (Supplementary Fig. 5a, iii-v). Notably, a density near residues Ser115 and Tyr116 appears conserved across all ATTR fibril structures determined to date (Fig. 5a-b, v-vi)^10-12,14-16^.

### Brain-derived and peripheral fibrils differ but share a conserved aggregation-driving region

Comparison of brain-derived and peripheral V30M fibrils reveals marked structural differences (Fig. 3). For this analysis, we selected heart-derived V30M fibrils as the representative peripheral structure with a closed gate fold^14^. The main differences include: (i) a continuous ordered core in brain-derived fibrils, compared with a segmented core in peripheral fibrils (Fig. 3a-b); (ii) a distinct orientation of the N-terminal region (Fig. 3c); and (iii) differences in the conformation of the gate region spanning Gly57–Gly67 (Fig. 3b). Comparison of brain-derived V30M and V30G fibrils further highlights pronounced differences in core size and organization, with V30G lacking large portions of the N- and C-terminal regions and adopting an alternative packing arrangement (Fig. 3a-b).

**Figure 3:**
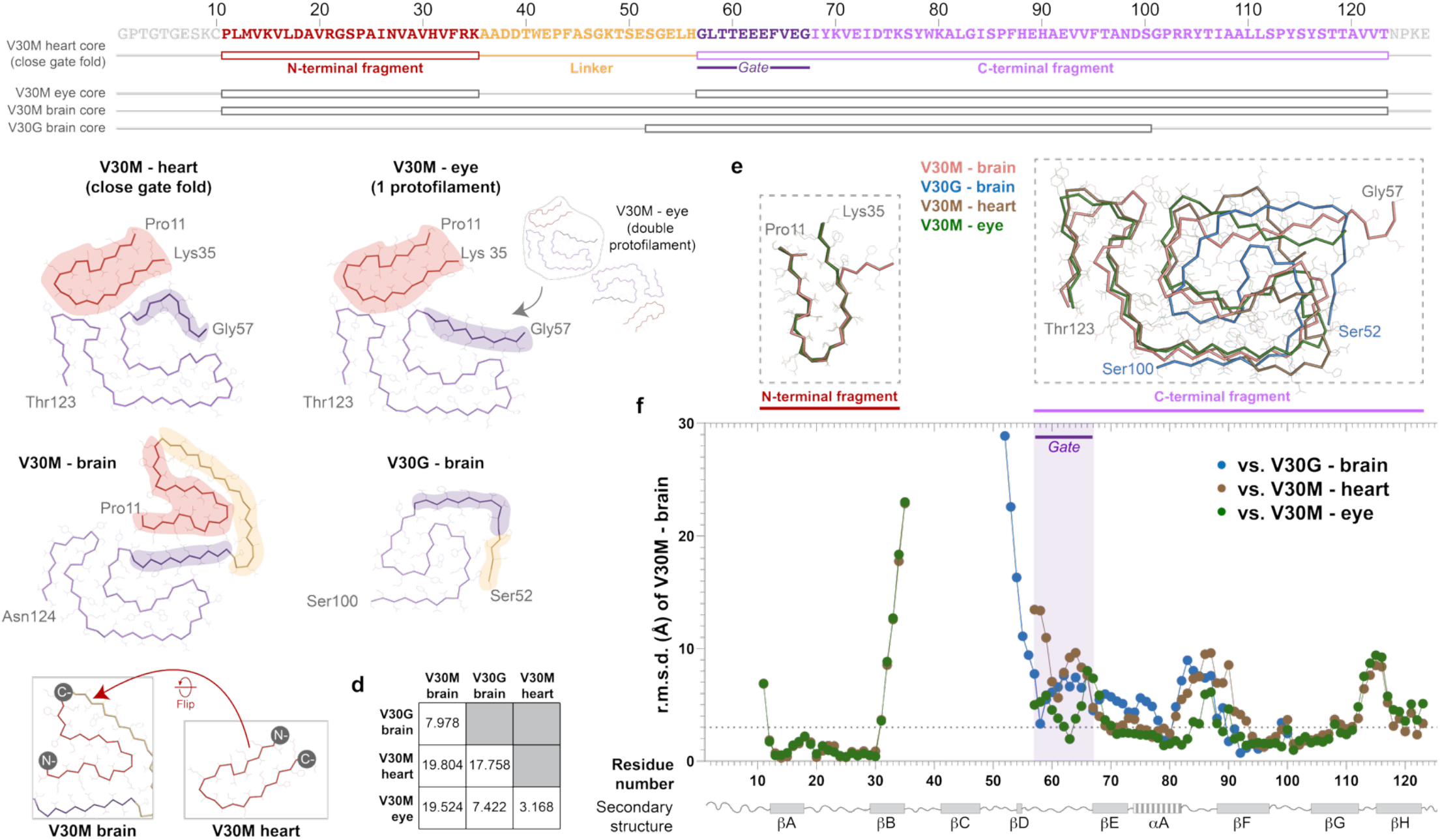
Structural comparison of ATTR fibrils. **a**, Schematic representation of transthyretin sequence organization highlighting regions present in fibril cores. The N-terminal fragment (red), linker (orange), gate region (purple), and C-terminal fragment (violet) defined from the V30M heart (PDB 9BZS) close gate fold are indicated. Colored bars denote residues included in the fibril cores of V30M heart (close gate), V30M eye (PDB 7OB4), V30M brain (PDB 13EV), and V30G brain (PDB 13EX). **b**, Top views of fibril core structures for V30M heart, V30M eye (single protofilament shown; double protofilament inset), V30M brain, and V30G brain, highlighting differences in core organization and connectivity. **c**, Structural comparison of the N-terminal fragment (Pro11–Lys35) between V30M brain and V30M heart fibrils, illustrating differences in orientation and packing. **d**, Matrix of pairwise root mean square deviation (r.m.s.d.) values for Cα alignment between fibril structures. **e**, Overlay of aligned fibril fragments. Left, alignment of the N-terminal fragment (Pro11–Lys35). Right, alignment of the C-terminal fragment (Gly57–Thr123), showing structural divergence across fibril types. Colors: V30M heart (brown), V30M eye (green), V30G brain (blue), and V30M brain (salmon). **f**, Residue-wise backbone r.m.s.d. between V30M brain fibrils and other fibril types (V30G brain, V30M heart, and V30M eye), plotted as a function of residue number. Shaded regions correspond to the N-terminal fragment (red), linker (orange), gate region (dark purple), and C-terminal fragment (light purple). Secondary structure elements from the native transthyretin tetramer are shown below.

These differences are supported by complementary computational analyses, including overall backbone comparisons by root mean square deviation, or r.m.s.d. (Fig. 3d), and side-chain packing differences quantified by the amyloid packing difference metric, or APD (Supplementary Fig. 6)^31^. Pairwise overall r.m.s.d. comparison of brain-derived V30M fibrils with other ATTR folds showed values above 7 Å, consistent with substantial structural divergence (Fig. 3e). APD analysis, which quantifies the percentage of residues involved in unique cross-β packing interactions or different side-chain orientations relative to the β-strands, further supports this conclusion. This analysis generates contact maps and APD values that allow clustering of amyloid folds^31^. In our dataset, brain-derived structures are structurally distant from one another and from peripheral ATTR fibrils, whereas peripheral ATTR fibrils cluster together with APD values below 25% (Supplementary Fig. 7). Together, these analyses demonstrate substantial structural divergence both across tissues and between variants within the brain.

Despite these pronounced differences, a subset of regions remains structurally conserved across all fibril structures, encompassing residues Leu12 to His31 and Phe95 to Ala109, when present (Fig. 3e-f). These stretches correspond to β-strands A, F, and G, and the loops A-B and F-G of the native transthyretin structure^19^ (Fig. 3g). Per-residue r.m.s.d. comparison of brain-derived V30M fibrils with the other folds supports this observation (Fig. 3f). Although the APD contacts maps show the formation of new contacts and residue orientations in the stretch from Leu12 to His31, the segment Phe95 to Ala109, corresponding to β-strands F and G, remain fairly conserved across fibril types (Supplementary Fig. 6a-d).

## Discussion

In this study we show that brain-derived transthyretin fibrils adopt structures that are distinct from those previously reported in peripheral tissues and the vitreous humor^10-18^. These differences are evident at multiple levels, including core organization and overall fold, and extend to both V30M and V30G variants. Notably, V30M brain fibrils exhibit a continuous N-to C-terminal fold, whereas peripheral fibrils are composed of disconnected segments, and V30G fibrils represent a more extreme departure with a reduced ordered core (Fig. 3a-b). Together, these findings demonstrate that transthyretin aggregation adopts multiple tissue-associated fibril folds.

Despite this divergence, we identify β-strand F as a structurally conserved element across fibril types, with important translational implications. β-strand F has been identified as a major driver of transthyretin aggregation and has been directly targeted for therapeutic development^19-25^. Notably, *coramitug*, a humanized monoclonal antibody currently being evaluated in a Phase 3 clinical trial for ATTR amyloidosis with cardiomyopathy, was designed to bind an epitope within this region^20,21^. The conservation of β-strand F across structurally distinct fibrils suggests that such antibodies may recognize a broad range of fibril conformations, even those present in the brain. Additionally, we developed TAD1, a structure-based probe designed to target β-strands F and H, that detects circulating aggregates of transthyretin in blood and is currently evaluated in a clinical trial (NCT07538518). This or similar β-strand F–targeting probes may enable detection of structurally diverse circulating aggregates across disease contexts, including those in the brain^32-34^. Conserved fibril regions may therefore support broadly applicable diagnostic and therapeutic approaches, whereas divergent regions may enable organ-selective targeting.

Our findings raise fundamental questions about where transthyretin aggregation is initiated and where fibril structure is defined. Previous work has shown that all peripheral organs within a given patient share a common fibril fold^12,14^, and that circulating aggregates are present prior to clinical symptoms^32-34^. These studies support a model in which fibril structure is established early and propagated through seeding^22^. Our data are consistent with this model in peripheral tissues but suggest a distinct scenario in the brain (Supplementary Figure 6e). In the brain, transthyretin is produced locally by the choroid plexus and secreted into the cerebrospinal fluid, and brain deposits are thought to derive predominantly from this local pool rather than from circulating protein^35^.The distinct biochemical composition of the cerebrospinal fluid relative to blood may therefore favor the formation of structurally divergent fibril conformations^36^.

Our data further indicate that primary sequence contributes to structural divergence. V30M and V30G fibrils, both formed within the brain, adopt markedly different folds, demonstrating that variation in sequence can influence fibril structure within a shared tissue environment (Fig. 2d, i). Similar sequence-dependent effects on fibril structures have been reported in tauopathies and synucleinopathies^37,38^. Amyloid structure is therefore not solely encoded by sequence but emerges from its interaction with tissue context.

ATTR amyloidosis provides a rare opportunity to disentangle the respective contributions of sequence and tissue context to amyloid structure. In most amyloid diseases, comparisons across tissues are confounded by differences in sequence, strain, or patient background, making it difficult to isolate the role of the local environment. In contrast, ATTR allows direct comparison of fibrils formed by the same protein sequence across multiple organs even from the same patient^12,14^. The relative structural conservation observed across peripheral ATTR fibrils establishes a baseline against which the divergence of brain-derived fibrils can be interpreted as an effect of local tissue context. These findings therefore extend beyond ATTR amyloidosis and provide a framework for understanding how local tissue context shapes amyloid structure.

## Conclusion

In summary, transthyretin fibrils formed in the brain adopt folds distinct from those observed in peripheral tissues, supporting a model in which sequence and tissue context jointly shape fibril structure. Despite differences in core organization, including a continuous structure in V30M fibrils and a reduced core in V30G fibrils, β-strand F remains conserved across fibril types, reflecting a balance between structural variation and conserved aggregation features with potential translational significance. More broadly, these findings provide direct evidence that local tissue context can shape amyloid fibril fold in human disease.

## Methods

### Patients and tissue material

Human brain tissue samples were obtained from the laboratory of late Dr. Merrill Benson at Indiana university and from the Dr. Ricardo Taipa from the Portuguese Brain Bank. The patient samples were fully anonymized prior to transfer and were therefore exempt from Institutional Review Board oversight at UT Southwestern medical Center. Informed consent for tissue collection and research use was obtained under the protocols governing the original studies conducted at Indiana University School of Medicine and at the Portuguese Brain Bank. No participants were recruited for the present study and no compensation was provided.

### Histology

Fresh-frozen tissue was thawed overnight at 4 °C and subsequently fixed in a 20-fold excess volume of 10% neutral buffered formalin for 48 h at room temperature with agitation. Samples were then transferred to 70% ethanol and processed for paraffin embedding following established protocols^39^. Sections from paraffin blocks were prepared for routine hematoxylin and eosin (H&E), Congo red, and Thioflavin-S staining. Regressive H&E staining was performed using a Sakura DRS-601 automated stainer (x, y, z configuration) with Leica Surgipath Selectech reagents, following methodology described in Sheehan’s textbook. Congo red–stained sections were counterstained with hematoxylin and evaluated under bright-field microscopy for amyloid deposition. Staining was carried out using 0.1% Congo red in alcoholic saline after sensitization in alkaline alcoholic saline (10% NaCl, 0.5% NaOH, 80% ethanol). Thioflavin-S–stained sections were examined for amyloid deposits exhibiting plaque-like and vascular patterns. Fluorescence was detected upon ultraviolet excitation (400–440 nm) with emission collected using a 470 nm long-pass filter. Staining followed an adapted protocol based on Guntern et al^40^. Briefly, paraffin sections were hydrated and subjected to sequential chemical treatments, including oxidation with potassium permanganate (0.25% w/v), bleaching with potassium metabisulfite (1%) and oxalic acid (1%), peroxidation with hydrogen peroxide (1%) in sodium hydroxide (2%), and acidification with acetic acid (0.25%), with water rinses between each step. Sections were then equilibrated in 50% ethanol and stained for 7 min in 0.004% Thioflavin-S in ethanol. Excess stain was removed by ethanol washes, followed by dehydration and clearing. Slides were mounted with Cytoseal 60 (Epredia), a non-fluorescent permanent mounting medium.

### Extraction of brain derived ATTR amyloid fibrils

ATTR amyloid fibrils were isolated from fresh-frozen human brain tissue following an adaptation of a previously established protocol with. Briefly, approximately 300 mg of frozen tissue was first thawed at room temperature and finely minced using a scalpel. The tissue pieces were then suspended in 0.5 mL of Tris-calcium buffer (20 mM Tris, 138 mM NaCl, 2 mM CaCl_2_, 0.1% NaN_3_, pH 8.0) and centrifuged at 13,000 × g for 5 minutes at 4 °C. The resulting pellet underwent four additional washes with the same Tris-calcium buffer. Following these washes, the pellet was incubated overnight at 37 °C with agitation at 700 rpm in 0.5 mL collagenase solution (5 mg mL^−1^ collagenase dissolved in Tris-calcium buffer) containing cOmplete EDTA-free protease inhibitor cocktail (Roche). After incubation, the mixture was centrifuged at 13,000 × g for 10 minutes at 4 °C. The pellet obtained was then resuspended in 0.5 mL of Tris– EDTA buffer (20 mM Tris, 140 mM NaCl, 10 mM EDTA, 0.1% NaN_3_, pH 8.0), followed by centrifugation at 13,000 × g for 5 minutes at 4°C. This washing step with Tris–EDTA buffer was repeated five more times. After completing these washes, fibrils were extracted by resuspending the pellet in 120 μL ice-cold water supplemented with 5 mM EDTA and centrifuging at 5,000 × g for 5 minutes at 4°C. The fibril extraction procedure was repeated twice more.

### Negative-stained transmission electron microscopy

The presence of ATTR amyloid fibril was confirmed by transmission electron microscopy as described. Briefly, a 3.5 μL fibril sample was spotted onto a freshly glow-discharged carbon film 300 mesh copper grid (Electron Microscopy Sciences), incubated for 1 min, and gently blotted onto a filter paper to remove the solution. 4 μL uranyl acetate was applied onto the grid and immediately removed. The grid was negatively stained with 4 µL of 2% uranyl acetate for 1 min and gently blotted to remove the solution. A FEI Tecnai G2 Spirit Biotwin electron microscope (Thermo Fisher Scientific) or JEOL 1400 Plus at an accelerating voltage of 120 kV was used to examine the specimens.

### Immunoblotting

Western blot was performed on the extracted fibrils. Briefly, 0.5 µg of fibrils were dissolved in 4X LDS sample buffer and boiled for 10 minutes at 95 °C. The samples were then loaded and run onto Bio-Rad gel system using a Tris-MOPS-SDS running buffer. After electrophoresis, the proteins were transferred to a 0.2 µm nitrocellulose membrane. The membrane was probed with an in-house antibody targeting C terminal of human transthyretin protein (1:1000). Horseradish peroxidase-conjugated goat anti-rabbit IgG (dilution 1:1000, Invitrogen) was used as the secondary antibody. The protein bands were visualized using Promega Chemiluminescent Substrate on Azure imager, following the manufacturer’s instructions.

### Mass spectrometry (MS) sample preparation, data acquisition and analysis

0.5 µg of extracted ATTR fibrils were prepared for tryptic MS analysis by boiling in tricine SDS buffer and running on a Novex™ 16% tris-tricine gel. The gel was stained with Coomassie dye, and the ATTR smear was excised for MS analysis. Samples were digested overnight with trypsin, then cleaned up with solid-phase extraction before injection onto a Q Exactive HF mass spectrometer coupled to an Ultimate 3000 RSLC-Nano liquid chromatography system. MS scans were acquired at 120,000 resolution, with up to 20 MS/MS spectra obtained using higher-energy collisional dissociation. Data were analyzed using Proteome Discoverer v3.0 SP1, with peptide identification against the human reviewed protein database from UniProt. The mass spectrometry proteomics data has been deposited to MassIVE (a member of ProteomeXchange)^41^ and can be accessed at ProteomeXchange Accession Number: PXD078077. Link for reviewers to download data: ftp://MSV000101740@massive-ftp.ucsd.edu.

### Cryo-EM sample preparation, data collection, and processing

Extracted fibril samples (3 µL) were applied to glow-discharged R 1.2/1.3, 300 mesh, Cu grids (Quantifoil), blotted with filter paper to remove excess sample, and plunge-frozen in liquid ethane using a Vitrobot Mark IV (FEI/Thermo Fisher Scientific). Cryo-EM samples were screened on Talos Arctica at the Cryo-Electron Microscopy Facility (CEMF) at The University of Texas Southwestern Medical Center (UTSW), and the final datasets were collected on a 300 keV Titan Krios microscope (FEI/Thermo Fisher Scientific) with ColdFEG/Falcon4i/Selectris operated at slit width of 10 eV at the Stanford-SLAC Cryo-EM Center (S^2^C^2^). Pixel size, frame rate, dose rate, final dose, and number of micrographs per sample are detailed in (Supplementary Table 2). The raw movie frames were gain-corrected, aligned, motion-corrected and dose-weighted using RELION’s own implemented motion correction program. Contrast transfer function (CTF) estimation was performed using CTFFIND 4.1^42^. All steps of helical reconstruction, three-dimensional (3D) refinement, and post-process were carried out using RELION 4.0^43,44^. The filaments were picked automatically using Topaz in RELION 4.0. Particles were extracted using a box size of 300 pixels with an inter-box distance of 3 asymmetrical units at helical rise of 4.8 Å. 2D classification of 300-pixel particles was used to estimate the fibril crossover distance. and to remove suboptimal segments. A featureless cylinder was used as an initial model. Fibril helix was assumed left-handed for 3D reconstruction and 3D auto refinements were performed. Subsequent 3D auto refinements with optimization of helical twist and rise were carried out once the estimated resolution of the map reached beyond 4.75 Å. 3D classifications without particle alignment were used to further remove suboptimal segments, and to separate different conformation as in the case of the double filaments. Particles potentially leading to the best reconstructed map were further chosen and run through additional 3D auto-refinements. CTF refinements, 3D auto refinements, and post-processing were repeated to obtain higher resolution. The final overall resolution was estimated from Fourier shell correlations at 0.143 threshold between two independently refined half-maps (Supplementary Fig. 1).

### Atomic model building and refinement

We used the automated machine-learning ModelAngelo approach with minor modifications to obtain an initial atomic model^45^. First, COOT v0.9.8.1 was used to build the peptide backbone of a single fibril layer featuring all alanine residues^46^. We then created a new density map of one fibril layer in ChimeraX v1.8 using the command ‘vol zone #1 near #2 range 3 new true’, where #1 refers to the post-processed fibril density map and #2 is the peptide backbone model^47^. This new map was then entered into ModelAngelo, both with and without the primary transthyretin sequence, to obtain the initial atomic models of ATTR fibrils. Using COOT, we made residue modifications and real-space refinements to finalize the model. Further refinement was carried out using ‘phenix.real_space_refine’ from PHENIX 1.20^48^. ChimeraX v1.8 was used for molecular graphics and structural analysis^49^. Model statistics are summarized in Supplementary Table 2.

### Stabilization energy calculation

The stabilization energy per residue was calculated by the sum of the products of the area buried for each atom and the corresponding atomic solvation parameters (Supplementary Fig. 3). The overall energy was calculated by the sum of energies of all residues, and different colors were assigned to each residue in the solvation energy map^50^.

### Amyloid Packing Difference calculation

The amyloid packing difference, or APD, was calculated using a previously published series of scripts^31^. Briefly, individual structures are analyzed for contacts between residues, generating a table of contacts. Then, the contacts are compared between two structures to analyze the percent of unique contacts between them, which is the APD. The brain-derived V30M and V30G structures, as well as the vitreous humor-derived (PDB 7OB4) and heart-derived (PDB 9BZS) V30M structures, were all analyzed pairwise for their APDs using the published contacts.py and compare.py scripts. We also performed the APD analysis using all available PDB structures found in the Amyloid Atlas (at the time of writing) for comparison to the brain-derived V30G and V30M fibril structures^50^. We used a total of 30 structures for our analysis, comprised of the two brain-derived structures and 28 out of 29 publicly available *ex-vivo* ATTR fibril structures. We excluded PDB structure 7Z40 since it was superseded by PDB structure 8ADE. We repeated the above analysis using Python scripts that act as wrappers for the published contacts.py and compare.py scripts but perform the analysis in a high-throughput manner and compare the 30 structures in a pairwise fashion. The scripts resulted in a matrix of pairwise APD values which we used to cluster the structures via the ‘average’ method in SciPy. The raw data for the APD analysis is available online via (available to reviewers upon request).

### Ordered residues comparison

We used a portion of the previously published CALYPSO pipeline to compare the ordered residues between the brain-derived fibril structures and the rest of the publicly available *ex-vivo* ATTR fibril structures via the Amyloid Atlas^50,51^. To do this, we included the brain-derived structures as local PDBs in a ‘Local’ directory before running the CALYPSO analysis, as directed by the CALYPSO protocol.

### Figure panels

All figure panels were created with Adobe Illustrator.

## Data availability

Mass spectrometry data have been deposited to MassIVE database (a member of ProteomeXchange) under accession code PXD078077. Graphed data is provided in the source data file. Cryo-EM maps have been deposited in the Electron Microscopy Data Bank under accession codes as following; V30G (EMD-77029), V30M (EMDB-77028). The atomic models of V30G and V30M are available at the Protein Data Bank under accession code 13EX and 13EV respectively. All data generated or analyzed during this study that support the findings are available within this published article and its supplementary data files.

## Acknowledgements

We thank the individuals and their families for donating brain tissue and to Indiana University and Portuguese Brain Bank for making it available for research. We thank late Dr. Merril Benson for his contributions to the field of amyloidosis. We thank Virender Singh and Preeti Singh for their help with manual picking of fibrils from micrographs and Yasmin Ahmed for her help with histology. We extend our appreciation to all members of the Structural Biology Laboratory (SBL) and the staff of the UTSW Cryo-Electron Microscopy Facility and Electron Microscopy Core Facility for their technical assistance and access to instrumentation. The SBL and CEMF at UT Southwestern are partially funded by the Cancer Prevention & Research Institute of Texas (CPRIT; grant RP220582). The authors thank the UTSW Proteomics Core for assistance with proteomics analysis and the UTSW Histopathology core for their assistance with histological experiments. We also acknowledge the national cryo-EM center at Stanford-SLAC (project CA172) for providing access to state-of-the-art instrumentation, technical support, and data collection services. Some cryo-EM data for this study were acquired at the Stanford-SLAC Cryo-EM Center (S2C2), which is funded by the National Institute of General Medical Sciences (1R24GM154186). The views expressed here are solely those of the authors and do not necessarily reflect the official policies of the National Institutes of Health. We are especially grateful to Lisa B. Dunn and Dr. Alexandre Cassago at S2C2 for their invaluable support and assistance throughout data collection. Some of this work was performed at the National Center for CryoEM Access and Training (NCCAT) and the Simons Electron Microscopy Center located at the New York Structural Biology Center, supported by the NIH Common Fund Transformative High Resolution Cryo-Electron Microscopy program (U24 GM129539, and NIGMS R24 GM154192) and by grants from the Simons Foundation (SF349247) and NY State Assembly. Computational analyses were made possible in part by the BioHPC high-performance computing facility at the Lyda Hill Department of Bioinformatics, UT Southwestern Medical Center (https://portal.biohpc.swmed.edu). We also acknowledge the use of ChatGPT (OpenAI, 2025) to assist with textual refinement and manuscript editing. The authors reviewed and verified all AI-assisted content. This work was supported by the American Heart Association, the National Institutes of Health, National Heart, Lung, and Blood Institute, the Welch Foundation, and the UT Southwestern Endowments

## Author contributions

L.S and S.A. conceptualized the study. S.A., L.S. and B.A.N. developed the methodology. S.A. and B.A.N. performed cryo-EM experiments and data analysis, with support from A.Z. P.B. performed computational analyses. S.A. and P.B. carried out negative-stain electron microscopy. C.L., B.E., C.K., R.T. and I.R. performed histological analyses, and R.T. provided patient samples. M.C.F.R. performed mass spectrometry analyses. R.P. conducted biochemical characterization, including immunoblotting. L.V. and M.M assisted with fibril extraction. L.S, S.A, B.A.N, and P.B contributed to the manuscript writing. All authors reviewed the writing. L.S secured funding.

