## Supplemental Figures and Table for "Transthyretin amyloid fibrils adopt distinct folds in the brain"

**Affiliations:**
*[1]Center for Alzheimer's and Neurodegenerative Diseases, Department of Biophysics, Peter O'Donnell Jr Brain Institute, University of Texas Southwestern Medical Center (UTSW), Dallas, TX, USA.*
*[2]AmyGo Solutions, Inc., Dallas, Texas, USA.*
*[3]Department of Internal Medicine, University of Texas Southwestern Medical Center (UTSW), Dallas, TX, USA.*
*[4]Portuguese Brain Bank, Neuropathology Unit, Department of Neurosciences, Centro Hospitalar Universitário do Porto, Largo Prof. Abel Salazar, 4099-001, Porto, Portugal.*
*[5]UMIB-Unit for Multidisciplinary Research in Biomedicine, ICBAS - School of Medicine and Biomedical Sciences, University of Porto, Porto, Portugal.*
*[6]Department of Pathology, University of Texas Southwestern Medical Center (UTSW), Dallas, TX, USA.*
*[7]Department of Ophthalmology, University of Texas Southwestern Medical Center (UTSW), Dallas, TX, USA.*

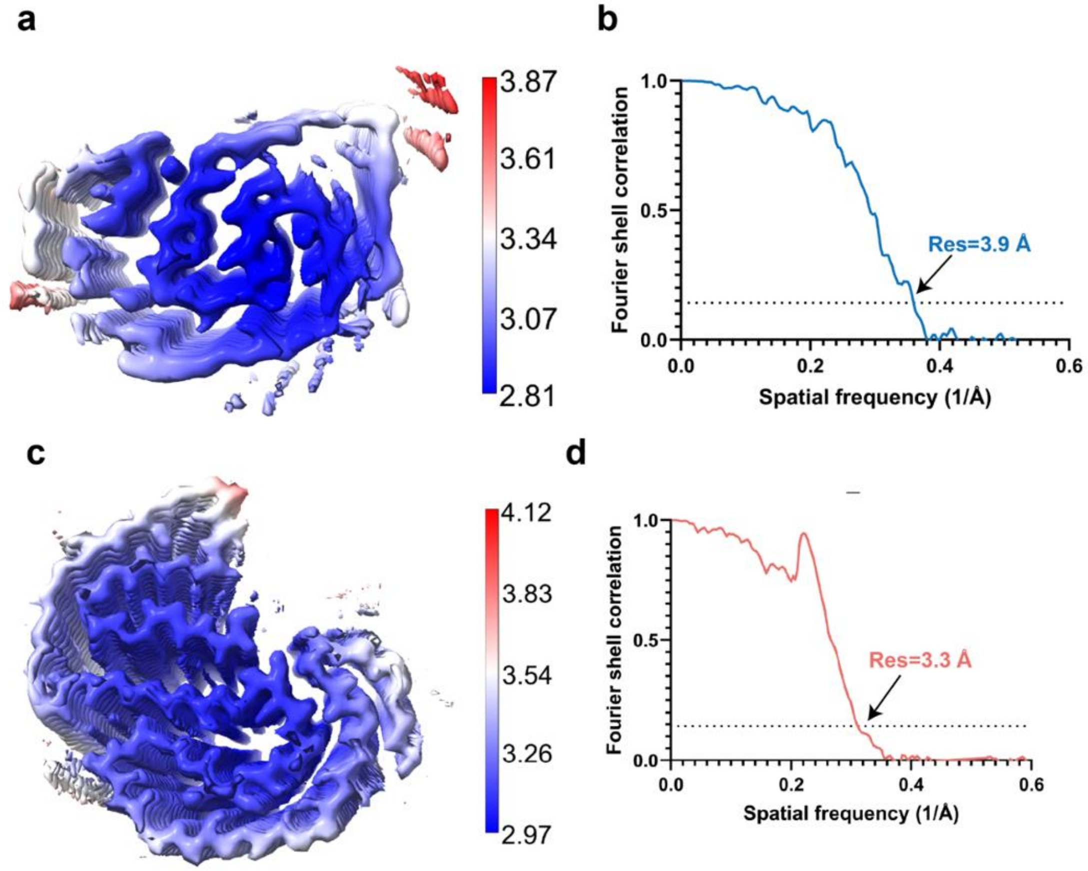


**Supplementary Fig. 1. Resolution assessment of the cryo-EM structures of V30G and V30M fibrils from the brain. a, c** Local resolution map of the V30G and V30M fibril structures, highlighting variations in resolution across different regions of the density map. **b, d.** Fourier shell correlation (FSC) curve between two independently refined half-maps, illustrating the overall resolution of the reconstruction for V30G and V30M fibril structures

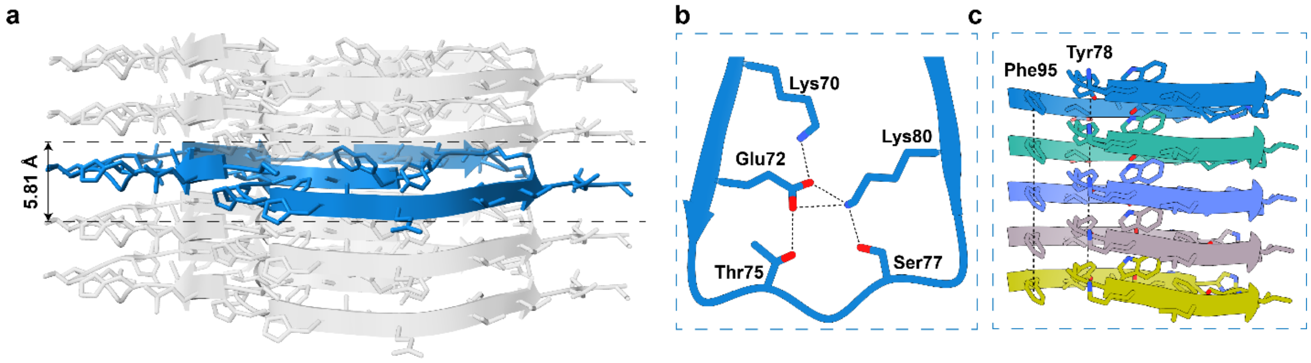


**Supplementary Fig. 2. Structural features stabilizing the ATTRv-V30G fibril architecture. a,** Five-layer representation of the ATTRv-V30G fibril model highlighting the in-register β-sheet arrangement along the fibril axis and a subunit height variation of 5.81Å. **b**. Intralayer salt bridges between Glu72–Lys70 and Glu72–Lys80, and hydrogen bonds between Thr75–Glu72 and Ser77–Lys80. **c**. Representative π–π stacking interactions are observed in Tyr78 and Phe95 residues across adjacent layers, contributing to further stabilization of the fibril structure. All interactions are noted as black dashed lines. Hydrogen bonding distance was between 2.5 Å and 3.1 Å. π-π stacking distance was between 4.8 Å and 4.9 Å.

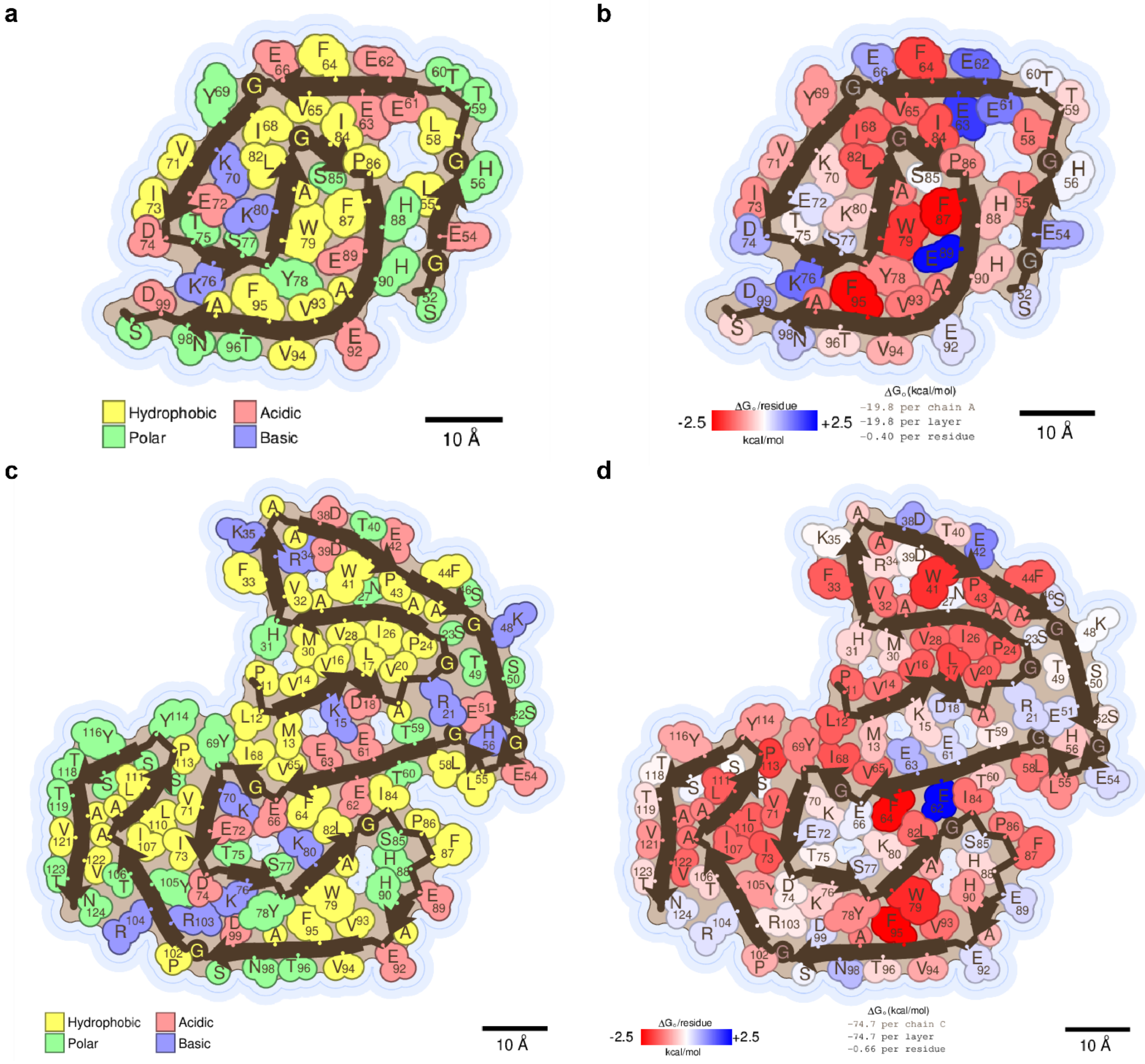


**Supplementary Fig.3. Composition and stability of the ATTRv brain fibril structures. a, c.** Schematic view of V30G (a) and V30M (c) fibrils showing residue composition where residues are color coded by amino acid category, as labeled. **b, d**. Representation of V30G (b) and V30M (d) fibril core depicting stabilizing residues. Strongly stabilizing side chains are colored red, and destabilizing side chains are colored blue.

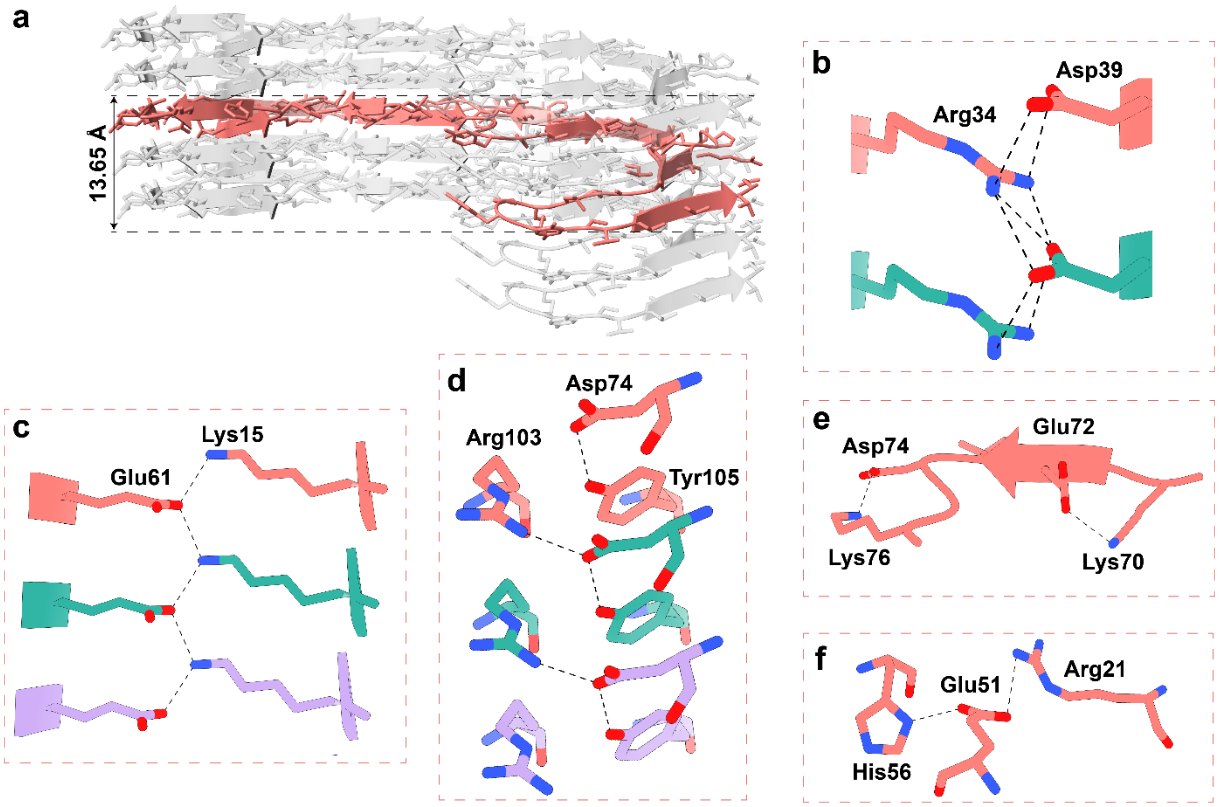


**Supplementary Fig.4. Additional interactions stabilizing the ATTRv-V30M brain fibril structure. a.** Five-layer representation of the ATTRv-V30M fibril model highlighting the in-register β-sheet arrangement along the fibril axis and a subunit height variation of 13.65 Å **b-d.** Representative interlayer salt bridge in V30M between Arg34 and Asp39 (b), Glu61 and Lys15 (c) and between Arg103 and Asp74, with an additional intralayer hydrogen bond between Asp74 and Tyr105 (d). **e-f.** Intralayer salt bridges between Asp74 and Lys76, Glu72 and Lys70 (e) and Glu51 and Arg 21 (f). All interactions are noted as black dashed lines. Hydrogen bonding distance was between 2.5 Å and 2.9 Å.

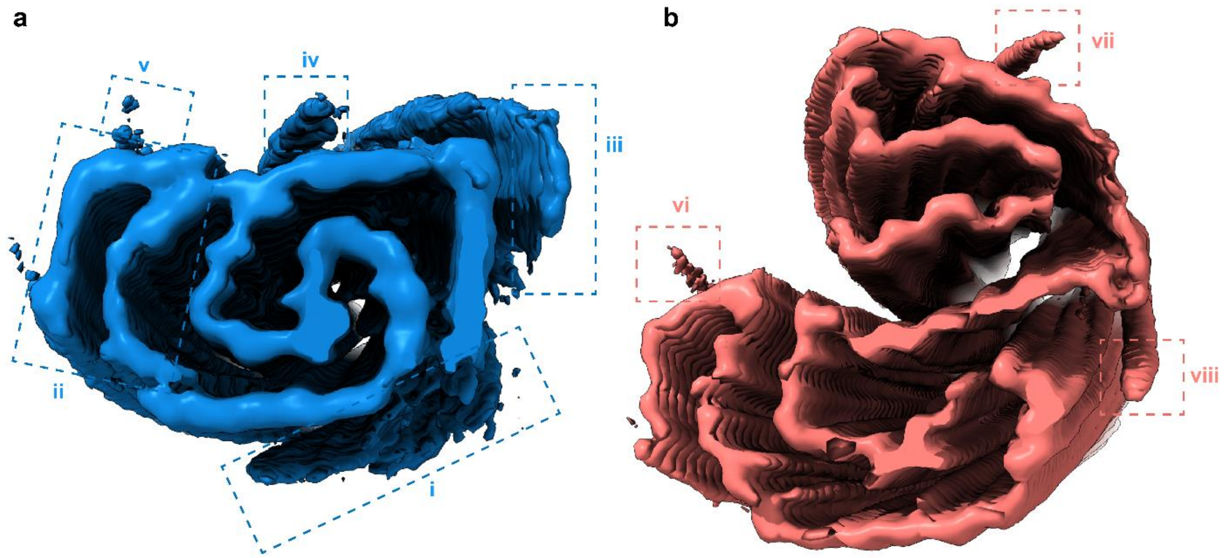


**Supplementary Fig.5. Unassigned extra densities in V30G and V30M fibril maps. a, b.** Cryo-EM density map of V30G (a) and V30M (b) fibrils highlighting unassigned densities (dashed boxes). These features are visible in the reconstructed map but could not be reliably modeled.

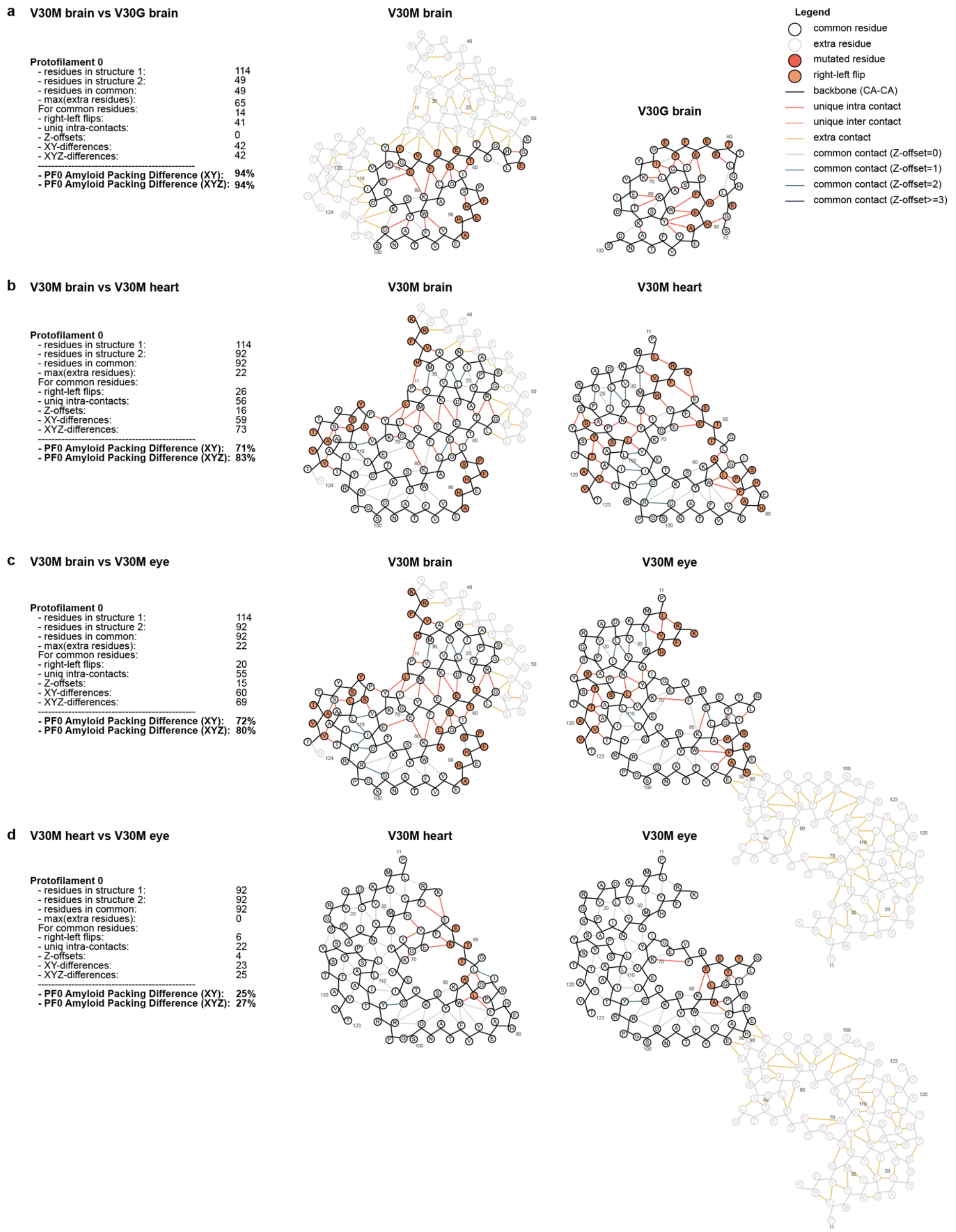


**Supplementary Fig. 6. Structural comparison and classification of *ex vivo* ATTR fibrils using XY APD scores**. Schematic representations of the atomic structures of chain A from **a.** V30M brain (PDB 13EV) vs V30G brain (PDB 13EX) **b.** V30M brain vs V30M heart (PDB 9bzs). **c.** V30M brain vs V30M eye (PDB 7OB4). **d.** V30M heart (PDB 9BZS) vs V30M eye (PDB 7OB4) as output by the compare.py script. The main chain is shown with black lines; residues with a side chain flip relative to their N-C-N planes in orange circles; contacts between residues with dark orange lines; common contacts without a Z-offset with grey lines; and common contacts with a Z-offset in teal green lines.

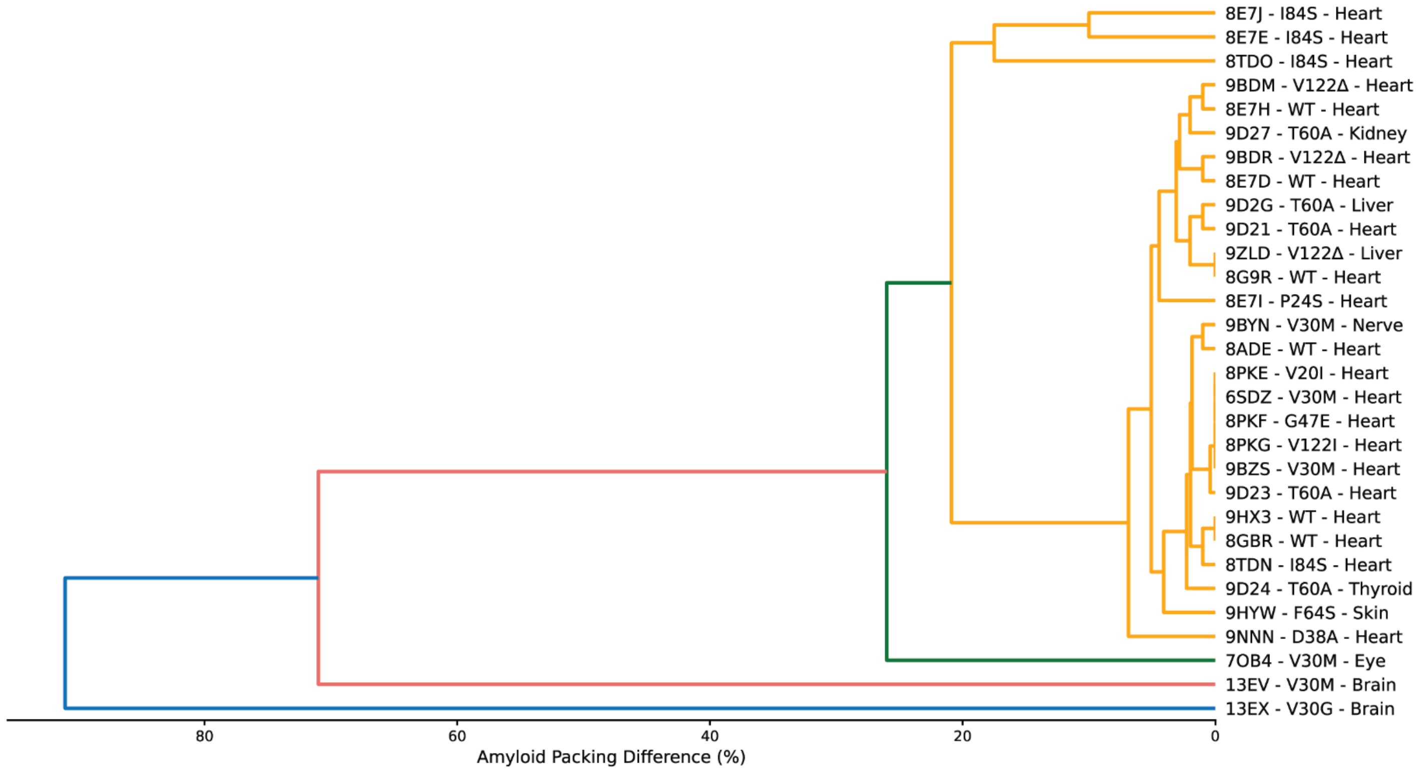


**Supplementary Fig. 7. Clustering of *ex vivo* ATTR fibril folds based on the APD.** PDB codes, mutation and source are shown for all clusters.

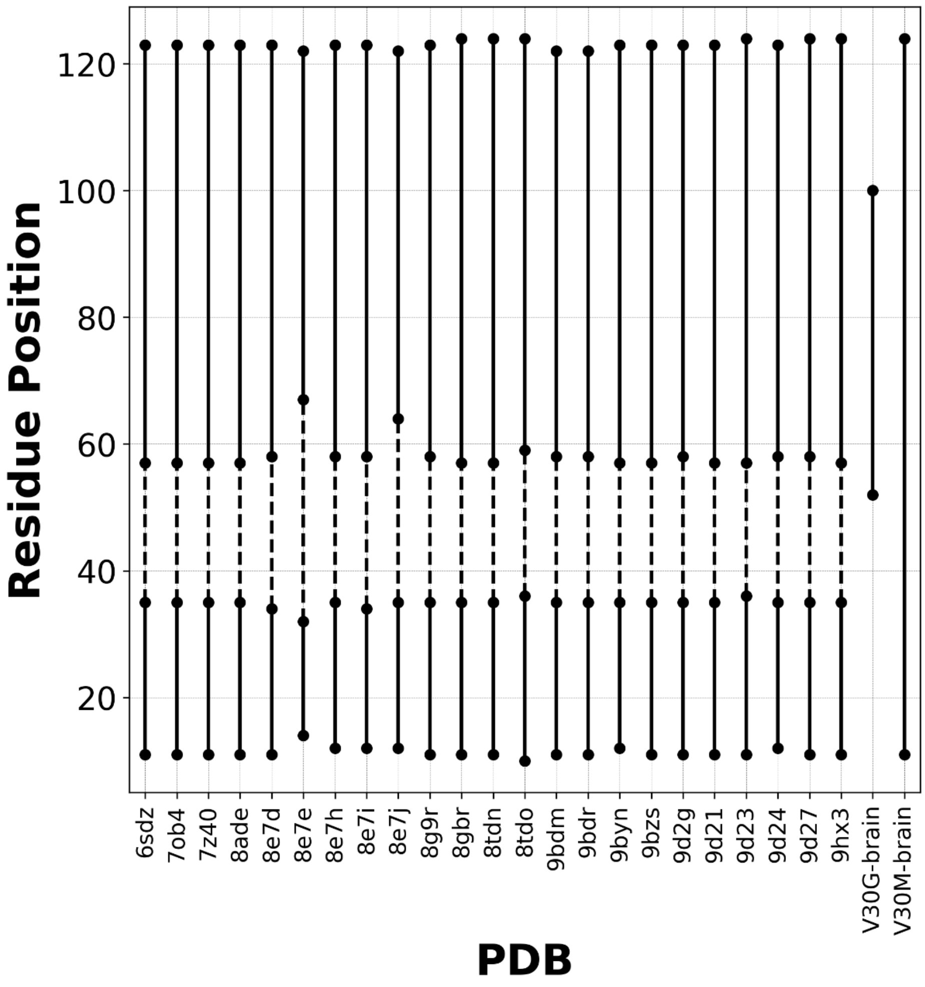


**Supplementary Fig 8.** Ordered residues in the core of *ex-vivo* ATTR fibrils.

**Supplementary table 1. Clinical history of patient samples used in the study.**

| Samples | Age at Onset (years) | Age at Death (years) | TTR Mutation |
|---|---|---|---|
| **Individual 1** | 45 | 45 | V30G |
| **Individual 2** | 34 | 63 | V30M |

**Supplementary Table 2. Cryo-EM data collection, refinement and validation statistics**

| | V30G (EMDB-77029) (PDB 13EX) | V30M (EMDB-77028) (PDB 13EV) |
|---|---|---|
| **Data collection and processing** | | |
| Magnification | 130,000 | |
| Voltage (kV) | 300 | 300 |
| Electron exposure (e–/Å$^2$) | 1 | 1 |
| Defocus range (μm) | -1.3 to -2.7 | -0.8 to -2.6 |
| Pixel size (Å) | 0.927 | 0.954 |
| Symmetry imposed | C1 | C1 |
| Helical twist | -0.89 | -1.04 |
| Helical rise | 4.85 | 4.75 |
| Initial particle images (no.) | 84,383 | 75,977 |
| Final particle images (no.) | 51,195 | 28,125 |
| Map resolution (Å) | 3.9 | 3.4 |
| FSC threshold (0.143) | | |
| Map resolution range (Å) | N/A | 2.97 – 3.83 |
| **Refinement** | | |
| Initial model used (PDB code) | Ab initio | Ab Initio |
| Model resolution (Å) | 3.4 | 3.0 |
| FSC threshold (0.143) | | |
| Map sharpening *B* factor (Å$^2$) | -37.72 | -57.63 |
| Model composition | | |
| Non-hydrogen atoms | 1950 | 4420 |
| Protein residues | 245 | 570 |
| *B* factors (Å$^2$) | | |
| Protein | 31.85 | 70.81 |
| R.m.s. deviations | | |
| Bond lengths (Å) | 0.004 | 0.005 |
| Bond angles (°) | 0.646 | 0.590 |
| Validation | | |
| MolProbity score | 2.07 | 1.72 |
| Clashscore | 13.87 | 9.18 |
| Poor rotamers (%) | 0 | 0 |
| Ramachandran plot | | |
| Favored (%) | 93.62 | 96.43 |
| Allowed (%) | 6.38 | 3.57 |
| Disallowed (%) | 0 | 0 |
